# Transcriptome analysis of an albino mutant in *Haworthia cooperi* var. *pilifera*

**DOI:** 10.1101/2020.11.24.396358

**Authors:** Maofei Ren, Yan Zhang, Hanbing Xu, Qingsong Zhu, Zhiyong Wang, Songhu Liu, Peiling Li, Benguo Liang

## Abstract

Photosynthetic organisms appear green due to the accumulation of chlorophyll (Chl) pigments in their chloroplasts. Although the genes encoding key enzymes related to Chl biosynthesis have been well characterized in herbaceous plants, such as rice, Arabidopsis and maize, white leaf mutants have not yet been fully studied in succulent plants. In this work, we explored the molecular mechanism of leaf color formation in an albino mutant (HUA) of *Haworthia cooperi* var. *pilifera*. We investigated the differentially expressed genes (DEGs) between HUA and control plants (wild type, LV) by transcriptome sequencing. Approximately 2,586 genes (1,996 downregulated and 590 upregulated) were found to be differentially expressed in HUA compared with LV using a threshold of ratio change ≥ 2 and false discovery rate (FDR) ≤0.05. GO analysis predicted that these DEGs participate in 12 cellular component, 20 biological process and 13 molecular function terms. Among the DEGs were well-recognized genes associated with chloroplast division and the biosynthesis of plant pigments, including chlorophyll, carotenoids and anthocyanin, as well as various transcription factor families. Overall, these results can help confirm the molecular regulatory mechanisms controlling leaf pigmentation and provide a comprehensive resource for breeding colorful leaf phenotypes in succulent plants.

## INTRODUCTION

Leaf-color mutants have recently received considerable attention as ideal materials for studying physiological processes in plants. A large number of such mutants have been identified in various plants, such as Arabidopsis (*Arabidopsis thaliana*)(Meier et al. 2011), rice (*Oryza sativa* L.)(Ki-Hong et al. 2003), soybean (*Glycine max* L.), maize (*Zea mays* L.)(Guan et al. 2016), and so on. There have been major advancements in understanding the genetic mechanisms of leaf-color mutations, and many related genes have been identified. Genes for chloroplast development and division, chlorophyll biosynthesis, carotenoid biosynthesis and anthocyanin synthesis have been identified in many plants, as well as transcription factors associated with leaf color. For example, GIK transcription factors coordinate the expression of the photosynthetic apparatus and are required for chloroplast development in Arabidopsis(Waters et al. 2009). ARC5, a cytosolic dynamin-like protein from plants, is part of the chloroplast division machinery(Gao et al. 2003). Mg-chelatase, a key enzyme for chlorophyll synthesis, comprises three subunits (ChlH, ChlD and ChlI) and catalyzes the insertion of Mg^2+^ into protoporphyrin IX, the last common intermediate precursor in both chlorophyll and heme biosynthesis, to produce Mg-protoporphyrin IX(Zhang et al. 2006). Many genes encode enzymes and transport proteins related to the biosynthesis of chlorophyll, carotenoids and anthocyanin, which have important roles in plants, such as protoporphyrinogen oxidase(Molina et al. 2010), zeta-carotene desaturase(Wong et al. 2004)and chalcone synthase(Ferrer et al. 2014). In addition, transcription factors such as MYB, bHLH and WD40 regulate the biosynthesis and transport of plant pigments by complex molecular mechanisms(Meng et al. 2014;Huq et al. 2004;Baudry et al. 2004).

*Haworthia* species are widely grown monocotyledonous desert succulents valued commercially as ornamentals(Campbel et al. 1995). *Haworthia cooperi* var. *pilifera*, a succulent plant from South Africa, is very popular in China and belongs to genus *Haworthia* and family Liliaceae. During the breeding of *Haworthia cooperi* var. *pilifera*, an albino mutant, which is quite different from the normal green plant, was obtained. Genomic information about this species is not yet available.

Next-generation sequencing (NGS) technologies have become a powerful approach for gene function annotation and gene expression analysis and have been used to identify numerous genes in many organisms14. (Sun et al. 2014). NGS contributes to molecular biology research in both model and nonmodel plants and is not dependent on an existing genomic sequence. Previous transcriptome sequencing analyses related to leaf color have focused on specific plants, such as birch (*Betula platyphylla* × *B. pendula*)(Gang et al. 2019) and tea (*Camellia sinensis* (L.) O. Kuntze)(Wang et al. 2014). A study of an *Acer palmatum* Thunb. golden-yellow mutant using transcriptome analysis revealed the pathway responsible for the golden-yellow mutation and uncovered novel genes related to leaf color(Li et al. 2015). The development of NGS provides us with more opportunities to discover novel genes and understand gene expression networks.

In this work, we performed transcriptome sequencing for the albino mutant (HUA) and green leaf *Haworthia cooperi* var. *pilifera* (LV) to characterize the gene expression in these two plants. We discovered candidate genes involved in chloroplast division and synthesis and transport of chlorophyll, carotenoids and anthocyanin. This study improves our understanding of the white-leaf phenotype in *Haworthia cooperi* var. *pilifera* and provides a basis for further studies of other succulent plants.

## MATERIALS AND METHODS

### Plant materials

*Haworthia cooperi* var. *pilifera*, a popular succulent cultivar, was used in this experiment. The seedlings were grown on MS medium with a thermoperiod of 25°C (day) and 22°C (night) and a photoperiod of 16 h. The leaves from healthy plants were harvested and stored at −80°C for subsequent RNA-Seq. Three biological replicates were performed for each group.

### RNA extraction, mRNA-seq library construction and sequencing

The total RNA of the two groups (control (LV) and albino mutant (HUA)) was extracted using a Plant RNA Kit (TaKaRa, Japan) following the manufacturer’s instructions. The quality and quantity of the total RNA were assessed at absorbance ratios of OD_260/280_ and OD_260/230_ and with 1% agarose gel electrophoresis. The replicates were mixed to produce 2 pooled samples for the sequencing analysis. After mRNA-seq libraries were generated using the VAHTS mRNA-seq v2 Library Prep Kit for Illumina (Vazyme, NR601) following the manufacturer’s recommendations, the library concentration was measured using the Qubit® RNA Assay Kit in Qubit® 3.0 for preliminary quantification. The insert size was assessed using the Agilent Bioanalysis 2100 system and found to be consistent with expectations; then, the qualified insert size was accurately quantified using qPCR with the StepOnePlus Real-Time PCR system (ABI, USA). The clustering of the index-coded samples was performed on a cBot Cluster Generation System (Illumina, USA) according to the manufacturer’s instructions. After cluster generation, the libraries were sequenced on an Illumina HiSeq X Ten platform with the 150 bp paired-end module.

### Sequence analysis, de novo assembly and annotation

The raw reads were cleaned by removing adaptor sequences, low-quality sequences (reads with ambiguous bases ‘N’), and reads with Q-value <20 bases. Then, we discarded reads with lengths less than 150 bp. Next, the remaining high-quality reads were *de novo* assembled into transcripts using the Trinity pipeline with the default settings. Then, the contigs were connected into transcript sequences to recover full-length transcripts across a broad range of expression levels, with the redundant duplicated reads removed. The longest transcript from the potential assembled alternative splicing transcript components was selected as the unigene of each set for subsequent analysis.

The unigenes were compared with public databases, and functional annotations were made based on similarity. The data used for comparison included the NCBI nonredundant proteins (NCBI NR), Gene Ontology (GO), Clusters of Orthologous Genes (COG), and Kyoto Encyclopedia of Genes and Genomes (KEGG) databases.

### Identification of differentially expressed genes (DEGs)

The DEGs in the HUA-LV comparison were defined using a FDR (false discovery rate) of ≤0.05 and an absolute value of log_2_ abundance ratio of ≥ 1 (twofold change). Then, the identified DEGs were subjected to GO functional analysis and KEGG pathway analysis.

## RESULTS

### Sequencing and de novo assembly

The main characteristics of the two RNA-seq libraries are summarized in Table 1. The number of raw reads per library ranged from ~4.21 to ~4.71 million. After filtering low-quality reads containing adaptors and low-quality sequences (defined as containing <90% confidently identified bases), we obtained 40,952,422 and 45,235,250 clean reads, which corresponded to 6,142,863,400 and 6,785,287,500 base pairs, respectively. The proportion of clean reads from each library was >96.04%. The transcriptomes of both the LV and HUA samples were represented by at least four million clean reads each, which allowed for quantitative analysis of gene transcription. The raw sequencing data have been submitted to the NCBI Sequence Read Archive (SRA) database with accession number SUB8069563.

**Table 1.** The statisics of sequencing

| Summary | LV | HUA |
| --- | --- | --- |
| Total raw reads | 42,199,192 | 47,096,827 |
| Total clean reads | 40,952,422 | 45,235,250 |
| Total clean nucleotides (bp) | 6,142,863,400 | 6,785,287,500 |
| Q20 (%) | 97.28 | 97.16 |
| Q30 (%) | 93.49 | 93.20 |
| N content (%) | 0.01 | 0.01 |
| GC content (%) | 49.08 | 48.75 |

Through *de novo* assembly, 84,163 unigenes with a mean length of 1,013 bp were obtained. The length of the assembled unigenes ranged from 200 to 15,563 bp with an N50 length of 1,668 bp (Table S1). The sequence length distribution of these unigenes is shown in Figure 1. Most unigenes were from 200-300 bp, while the longest unigene was 15,563 bp.

**Figure 1.**
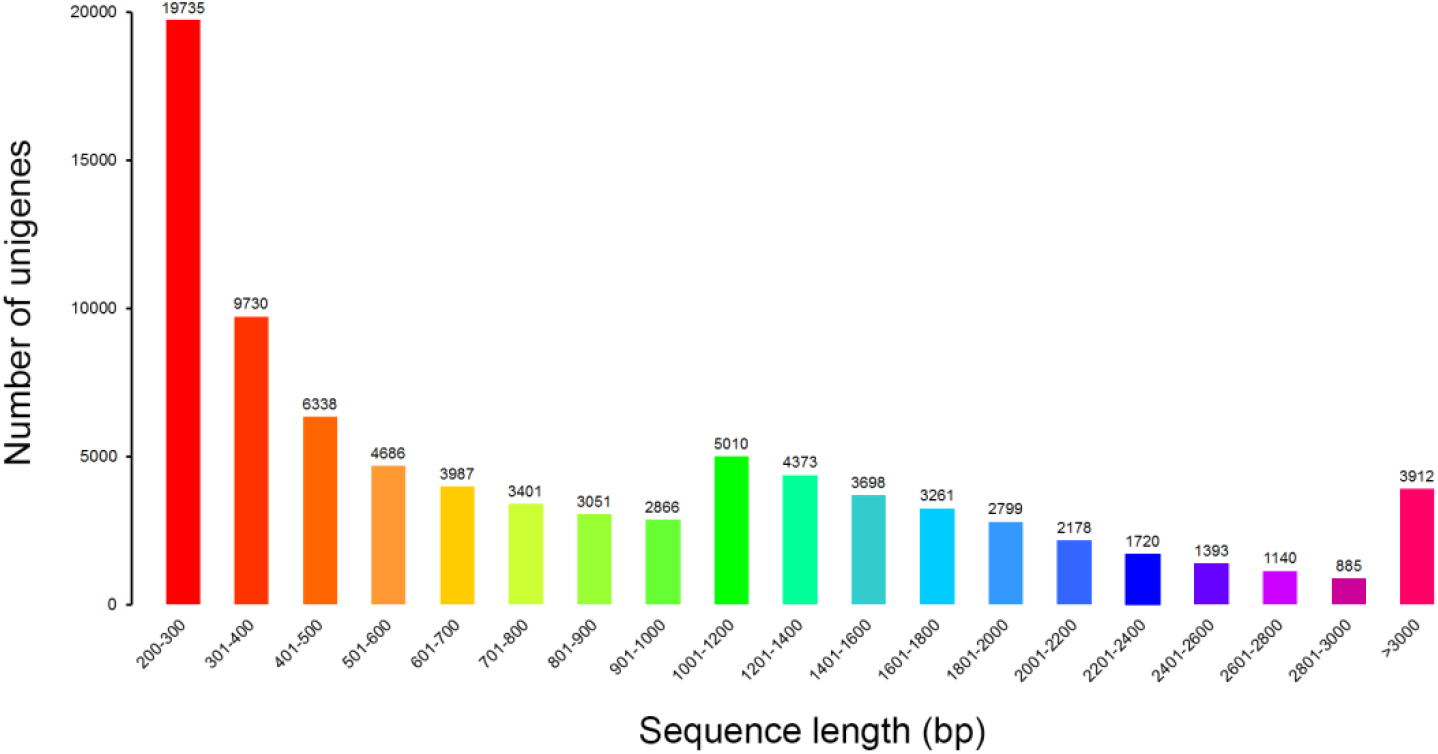
Length distribution of unigenes. The x-axis indicates sequence lengths from 1 to 20000 nt. The y-axis indicates the corresponding number of unigenes.

### Gene annotation and functional calculation

Alignment between the unigenes and public databases such as NR, NT, Swiss-Prot, KEGG, Clusters of Orthologous Genes (COG), and GO was performed, and the results with the best alignment were used to determine the sequence direction of each unigene. The number and percentage of unigenes aligned to each database are shown in Table S2. A total of 48,085 (57.13%), 36,210 (43.02%), 33,036 (39.25%), 30,353 (36.06%), 20,941 (24.88%) and 33,527 (39.84%) unigenes had significant hits in NR, NT, Swiss-Prot, KEGG, COG and GO, respectively.

For the E-value distribution, 58.26% of the unigenes had confidence levels of at least 1E-45 (Figure 2A). A total of 53.21% had alignment identities greater than 60% with existing proteins in the NR database (Figure 2B). To gain insight into the percentage of the 47,851 unigenes that showed similarity to genes in other plant species, we analyzed the species distribution of *Haworthia cooperi* var. *pilifera* in the NR database, in which 65.21% of the genes were annotated as having the highest similarity to genes from 8 species, the largest number among the databases examined (Figure 3). Among the various plants, the 12,378 unigenes had the highest number of unigenes with the greatest similarity to *Vitis vinifera* (25.87%), 8.93% to *Oryza sativa* Japonica Group, 5.83% to *Prunus persica*, 5.73% to *Ricinus communis*, and 5.04% to *Brachypodium distachyon*.

**Figure 2.**
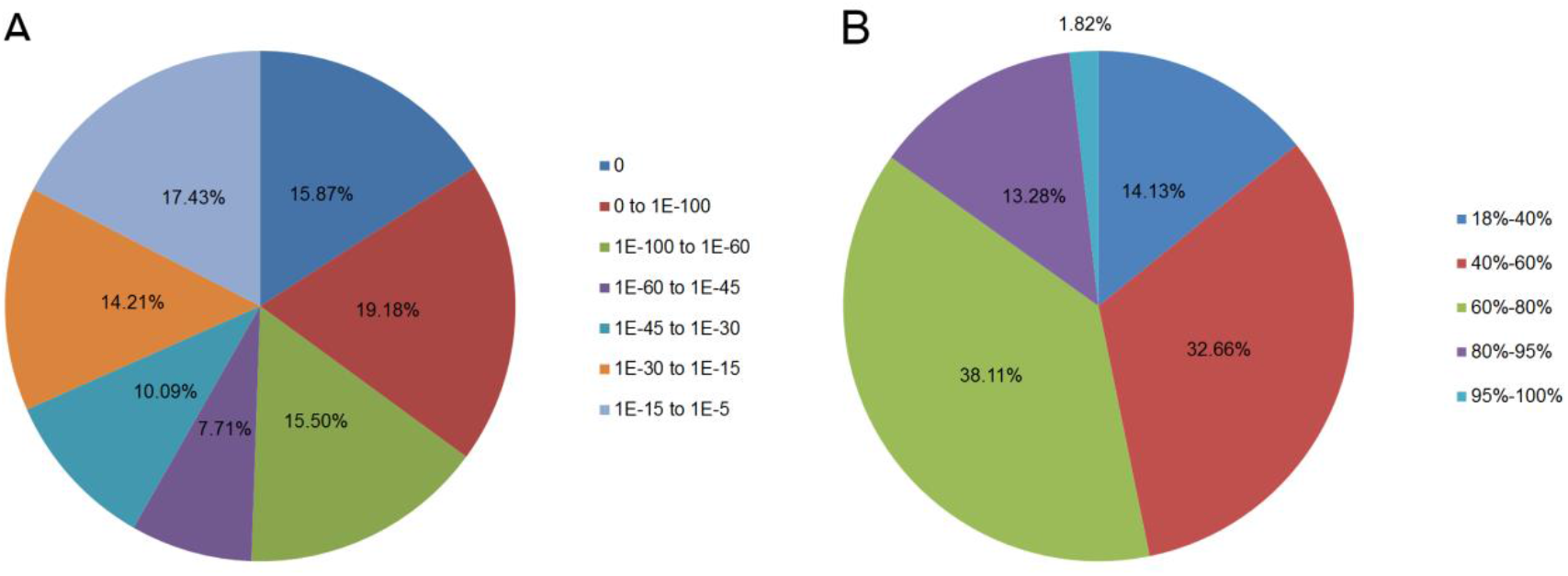
Homology search results for query sequences aligned by BLASTx to the NR database. (A) E-value distribution of unigene BLASTx hits in the NR database with an E-value threshold of 1.0 E-5. (B) Identity distribution of the top BLAST hits for each unigene.

**Figure 3.**
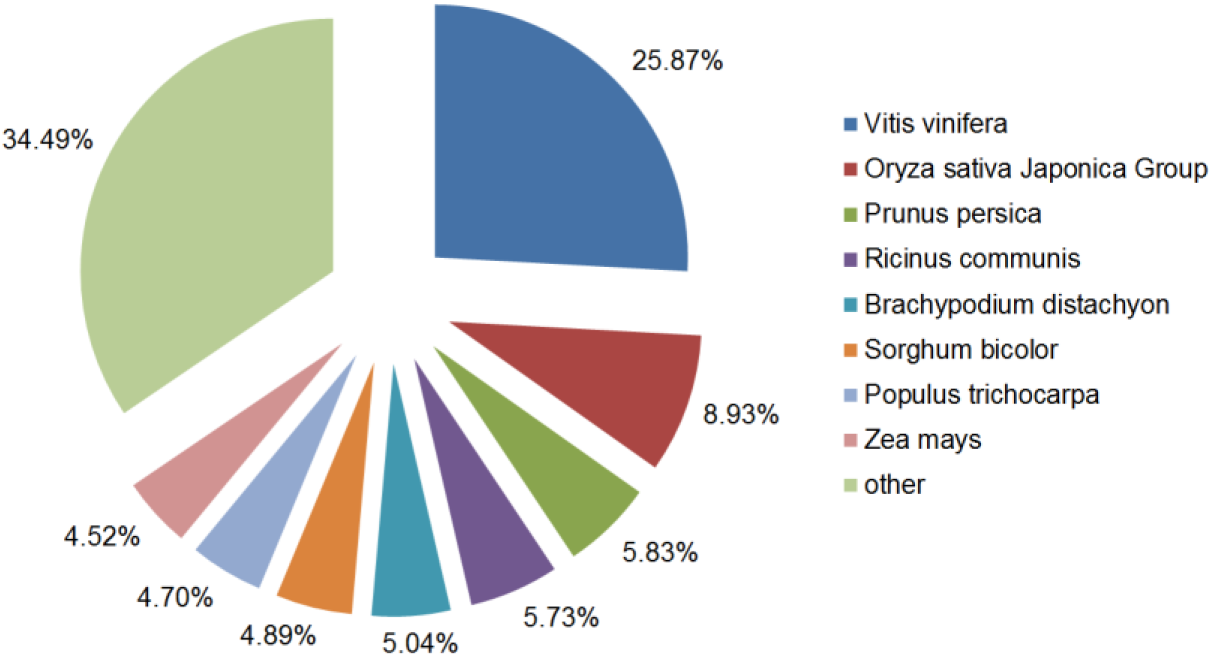
Percentage of the *Haworthia cooperi* var. *pilifera* unigenes showing similarity to genes in other plant species. The *Haworthia cooperi* var. *pilifera* unigenes were aligned by BLASTx to the NR database.

The GO analysis allowed the functional classification of 33,527 unigenes. As shown in Figure 4, the unigenes could be categorized into 56 functional groups on the terms of sequence homology. GO terms are grouped into three categories: biological process, cellular component and molecular function. In the biological process category, the largest number of unigenes were associated with ‘cellular process’ (59.55%), followed by ‘metabolic process’ (55.79%) and ‘single-organism process’ (33.67%). On the other hand, only a few genes were associated with ‘biological adhesion’ (0.12%), ‘locomotion’ (0.10%), and ‘cell killing’ (0.02%). The major cellular component terms in the GO classification were ‘cell’, ‘cell part’ and ‘organelle’, while the predominant molecular function terms were ‘binding’ and ‘catalytic activity’.

**Figure 4.**
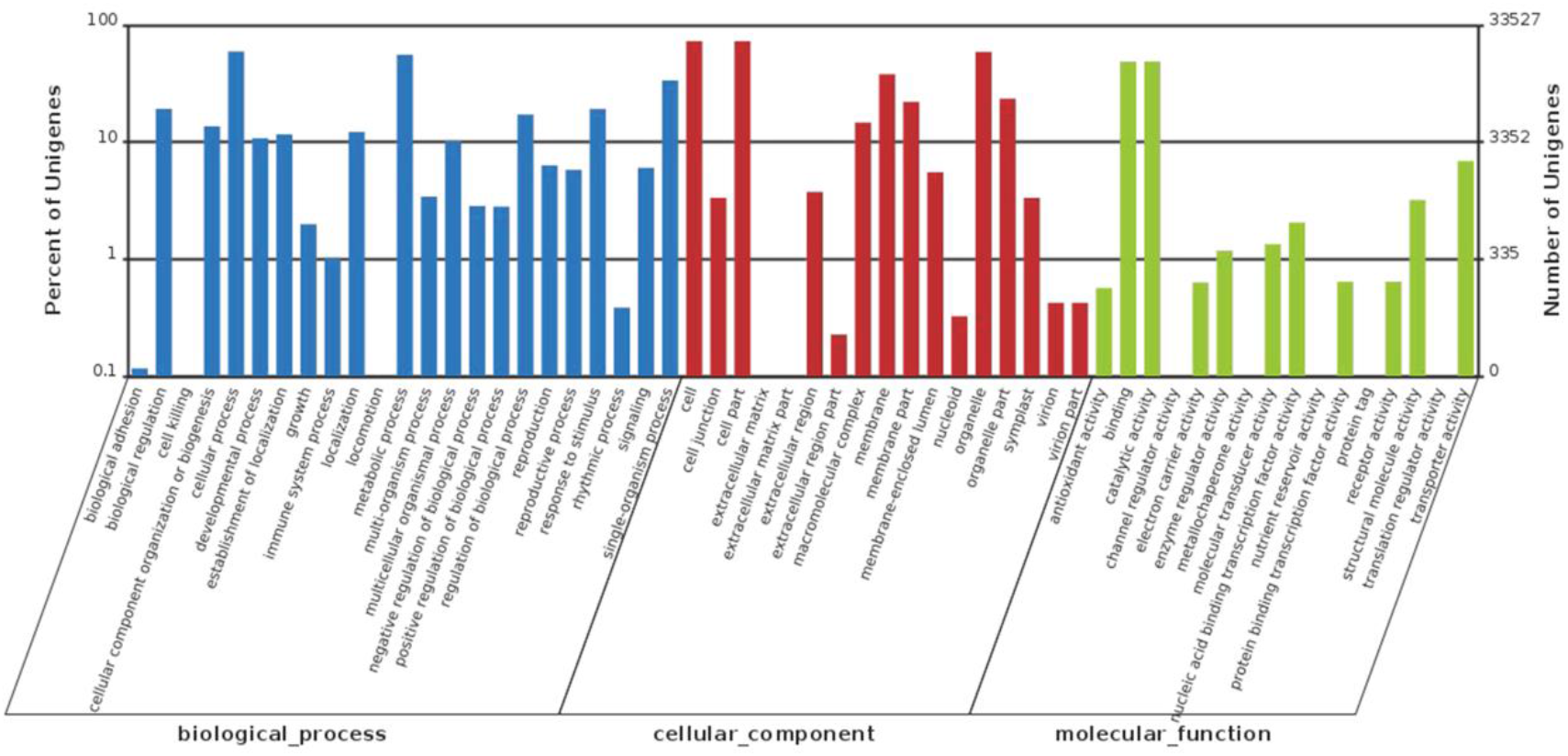
Distribution of Gene Ontology (GO) annotations of all unigenes from *Haworthia cooperi* var. *pilifera*. A total of 33,527 unigenes were assigned GO terms, and they were classified into three GO categories: biological process, cellular component and molecular function. The left y-axis indicates the percentage of unigenes in a specific category. The right y-axis indicates the number of unigenes in a category.

We further classified the functions of the unigenes by using the COG database. A total of 20,941 unigenes were assigned to 25 specific COG categories, which are listed in Figure 5. The largest group was ‘General function prediction only’ with 7,834 unigenes (37.41%), followed by ‘Transcription’ with 6,253 unigenes (29.86%), ‘Replication, recombination and repair’ with 4,778 unigenes (22.82%), ‘Posttranslational modification, protein turnover, chaperones’ with 3,888 unigenes (18.57%), and ‘Signal transduction mechanisms’ with 3,872 unigenes (18.49%). ‘Function unknown’ contained 3,529 unigenes (16.85%), ‘Translation, ribosomal structure and biogenesis’ contained 3,428 unigenes (16.37%), ‘Cell cycle control, cell division, chromosome partitioning’ contained 3,197 unigenes (15.27%), ‘Carbohydrate transport and metabolism’ contained 3,150 unigenes (15.04%), and ‘Cell wall/membrane/envelope biogenesis’ contained 2,712 unigenes (12.95%). The categories ‘Nuclear structure’ and ‘Extracellular structures’ represented the smallest groups.

**Figure 5.**
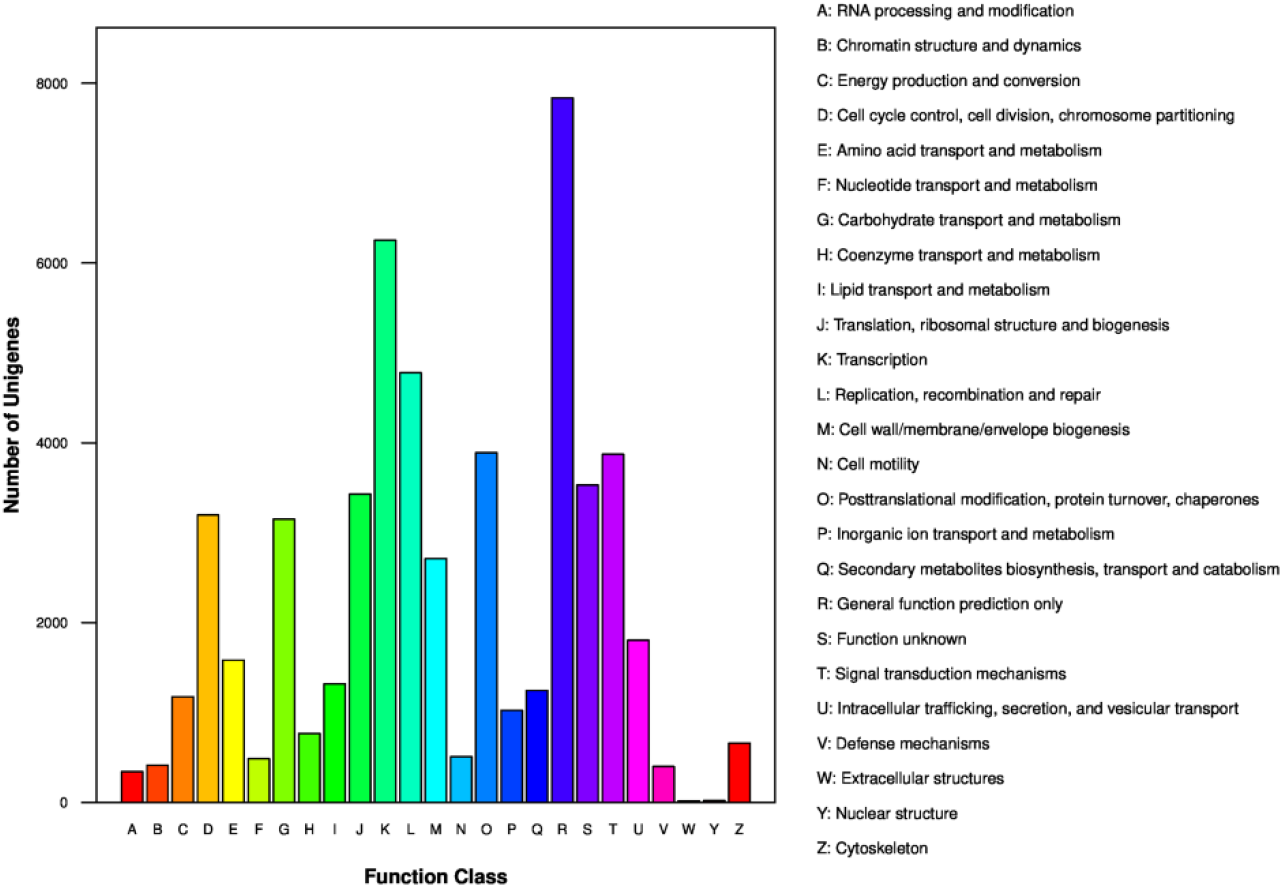
COG functional classifications of all unigenes from *Haworthia cooperi* var. *pilifera*. A total of 20,941 sequences had a COG classification among 25 categories.

KEGG was used as a higher-order functional annotation to understand the metabolic and biological pathways and functions of the gene products in the cells. We annotated and mapped 128 KEGG pathways for 30,353 unigenes (Figure 6). These pathways were distributed in five main categories: metabolism (97.33%), genetic information processing (35.37%), organismal systems (6.94%), cellular processes (9.22%) and environmental information processing (6.17%). The KEGG pathway analysis showed that the unigenes were mainly located in metabolic pathways (ko01100, 8,107 unigenes), biosynthesis of secondary metabolites (ko01110, 3,183 unigenes), endocytosis (ko04144, 2,079 unigenes) and glycerophospholipid metabolism (ko00564, 2,059 unigenes).

**Figure 6.**
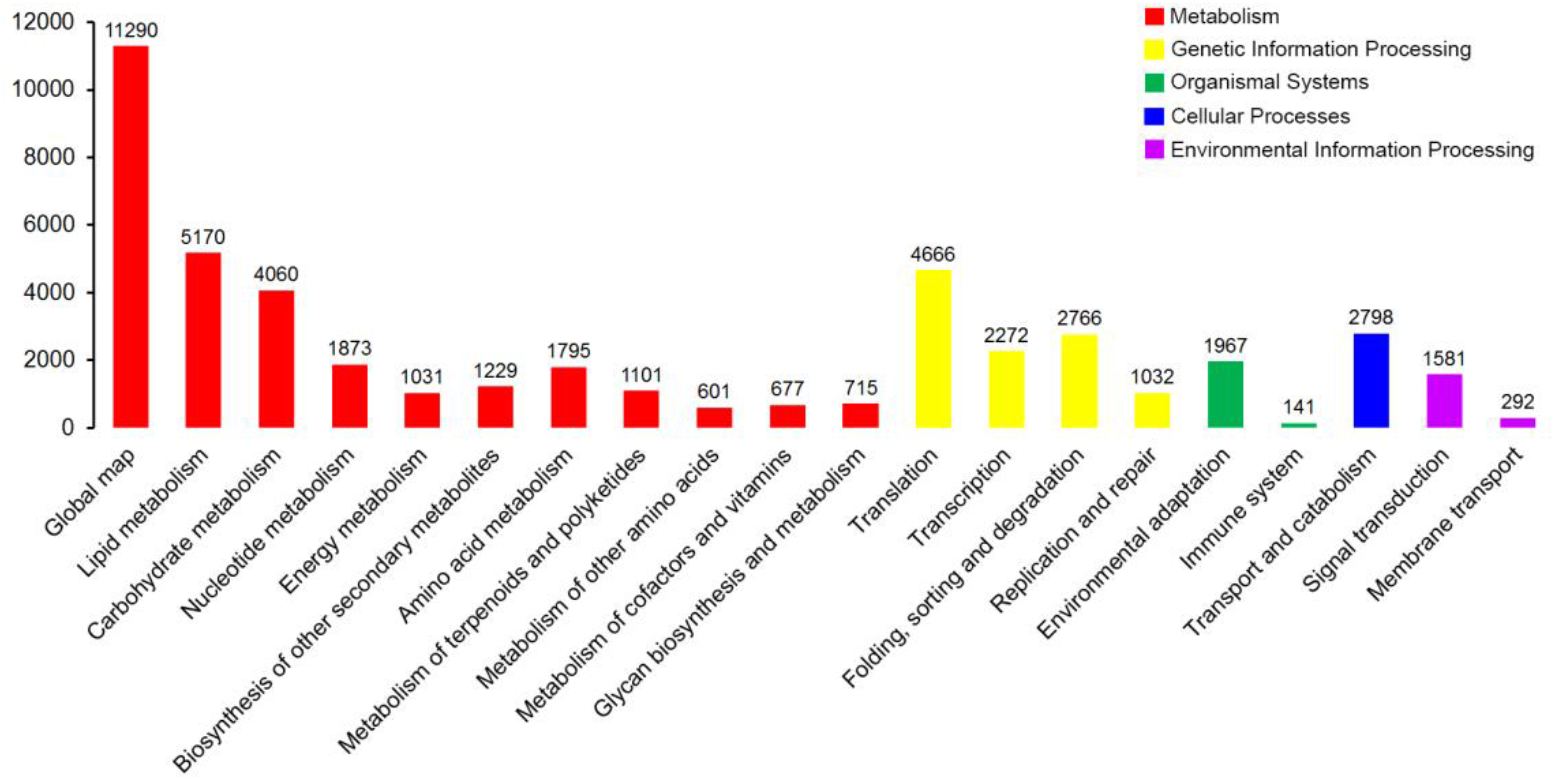
KEGG pathway assignments of the assembled unigenes. A total of 30,353 unigenes were mapped to 128 KEGG pathways belonging to five categories: metabolism (red), genetic information processing (yellow), organismal systems (green), cellular processes (blue) and environmental information processing (purple).

### Analysis of potential DEGs

Based on expression abundance (FPKM values), the gene expression levels of the two samples are shown in Figure 7. A higher number of downregulated than upregulated genes was observed in the HUA sample. In total, 590 DEGs showed upregulated expression, while 1,996 DEGs were downregulated (Table S3).

**Figure 7.**
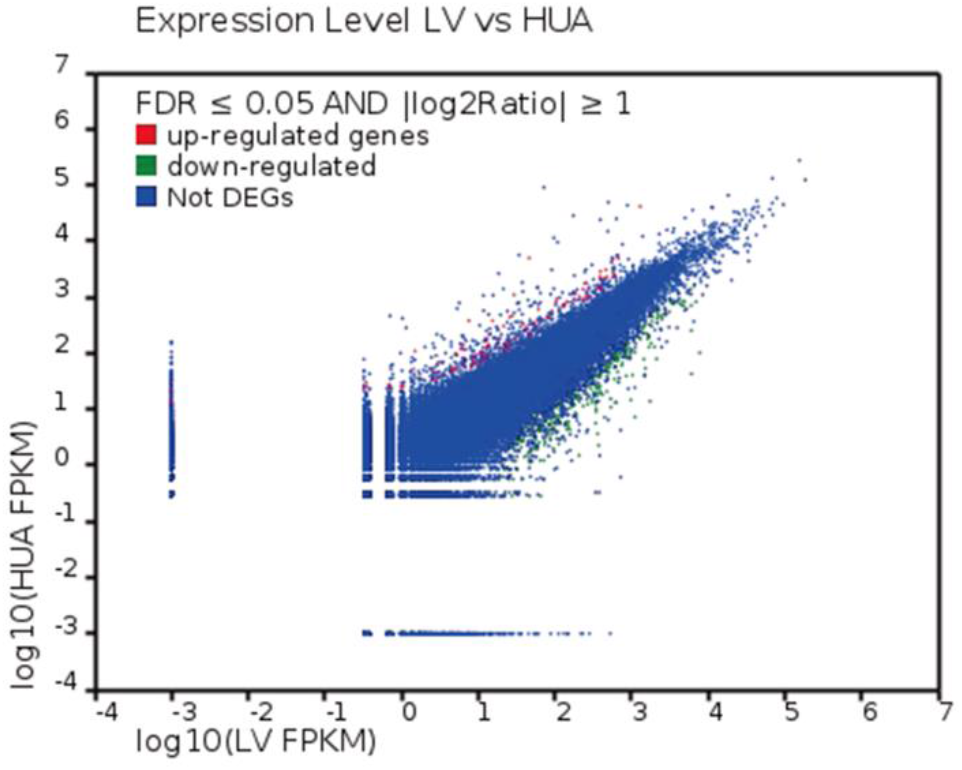
Gene expression levels between the two libraries (LV and HUA). Red dots represent transcripts more prevalent in the HUA library, green dots show those present at a lower frequency in HUA, while blue dots indicate transcripts that did not change significantly.

### GO and KEGG enrichment analyses of DEGs

GO annotation was used to determine the functions of the DEGs. The DEGs were divided into 45 functional terms in three categories: biological process, cellular component and molecular function **(Figure 8)**. Cell (846 DEGs) was the most common term, followed by cell part (842), cellular process (764), catalytic activity (735), metabolic process (727), single-organism process (609), organelle (609), binding (584) and membrane (541). The GO terms in all three ontologies with the highest enrichment of DEGs are shown in Table 2.

**Table 2.**
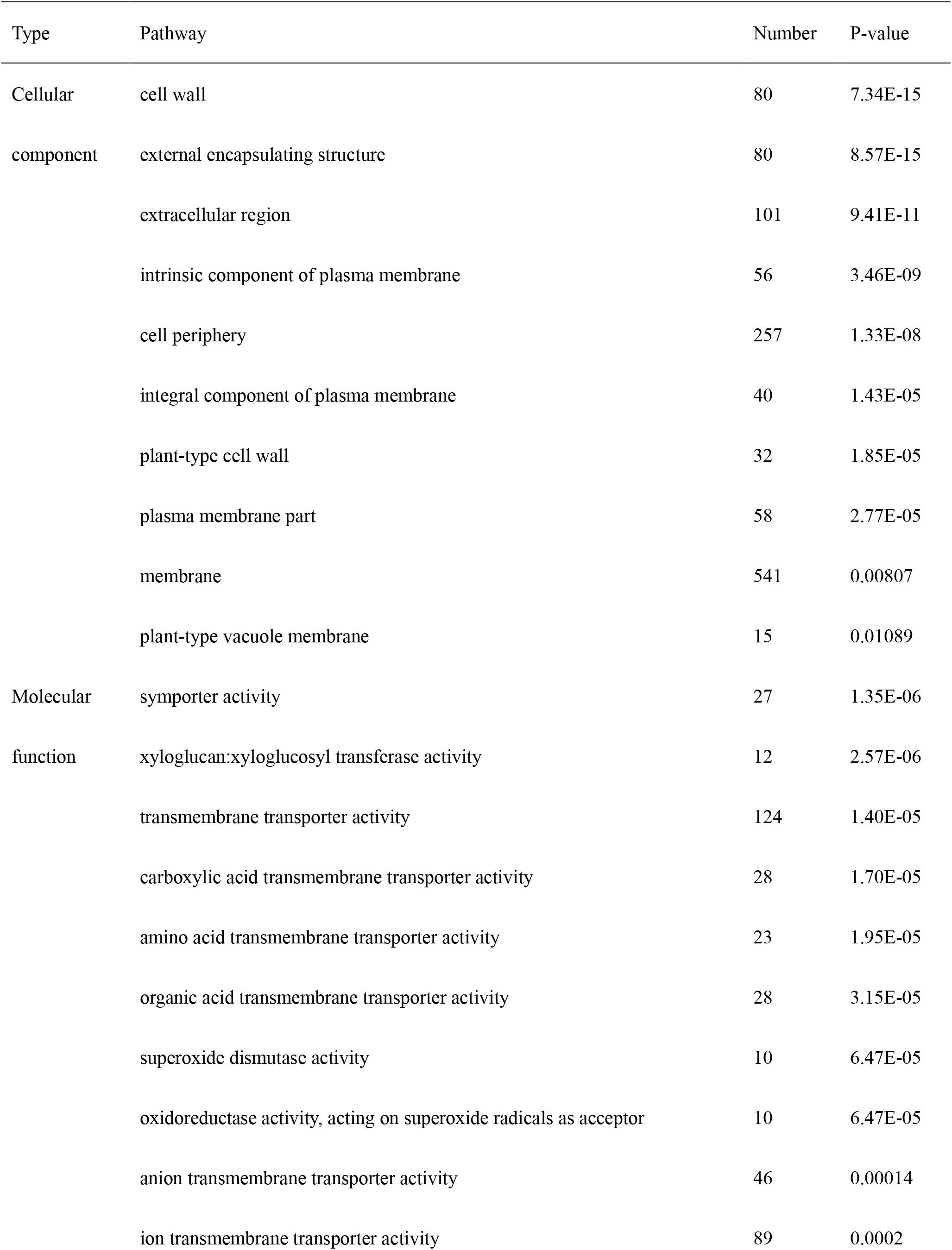
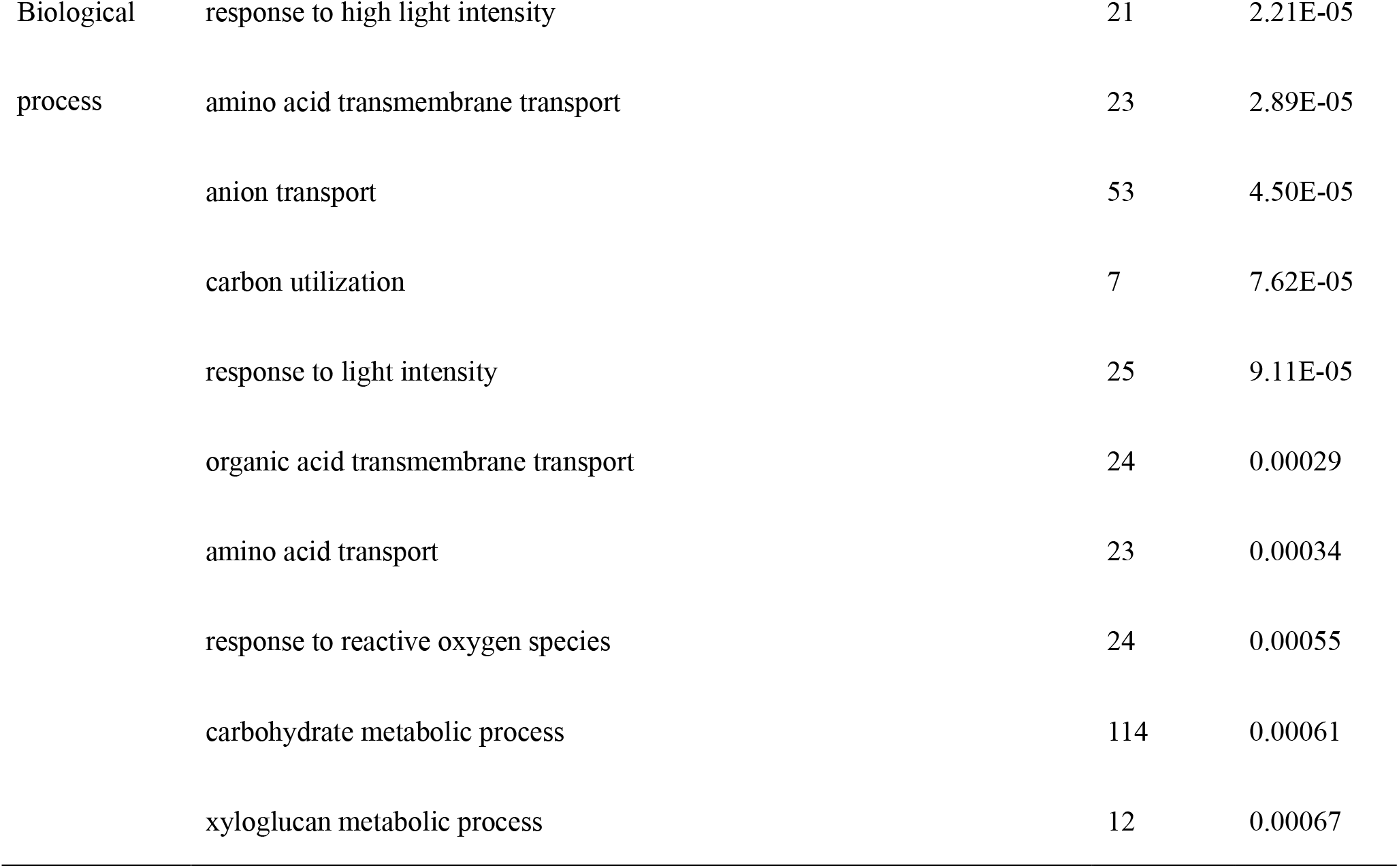
The most reliable top ten significantly enriched pathways of DEGs in Cellular component, molecular function and biological process

**Figure 8.**
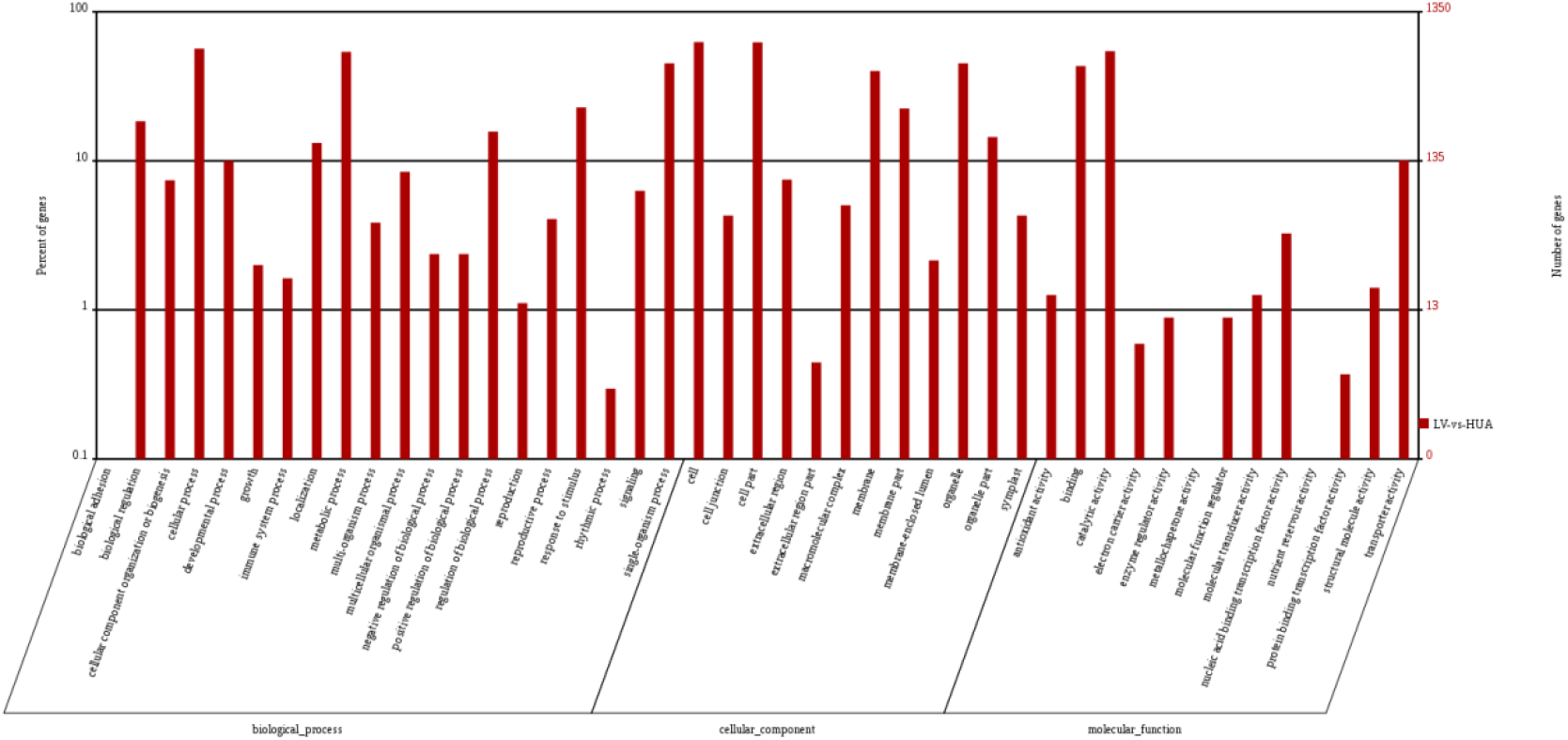
GO enrichment analysis of DEGs in LV vs HUA. Enrichment in the GO categories ‘biological process’, ‘cellular components’ and ‘molecular function’ categories is shown. Further classifications under each category are also listed.

To understand the biological functions of the DEGs, we also mapped these genes to pathways in the KEGG database. The DEGs were functionally assigned to 109 KEGG pathways, and the 20 pathways most enriched in DEGs are shown in Figure 9. Among the mapped pathways, 24 pathways were significantly enriched (P value≤ 0.01) (Table S4). ‘Metabolic pathways’ was the most enriched pathway (420 DEGs), followed by ‘Plant-pathogen interaction’ (149), ‘Glycerophospholipid metabolism’ (127), ‘Endocytosis’ (125), ‘Plant hormone signal transduction’ (124) and ‘Ether lipid metabolism’ (117).

**Figure 9.**
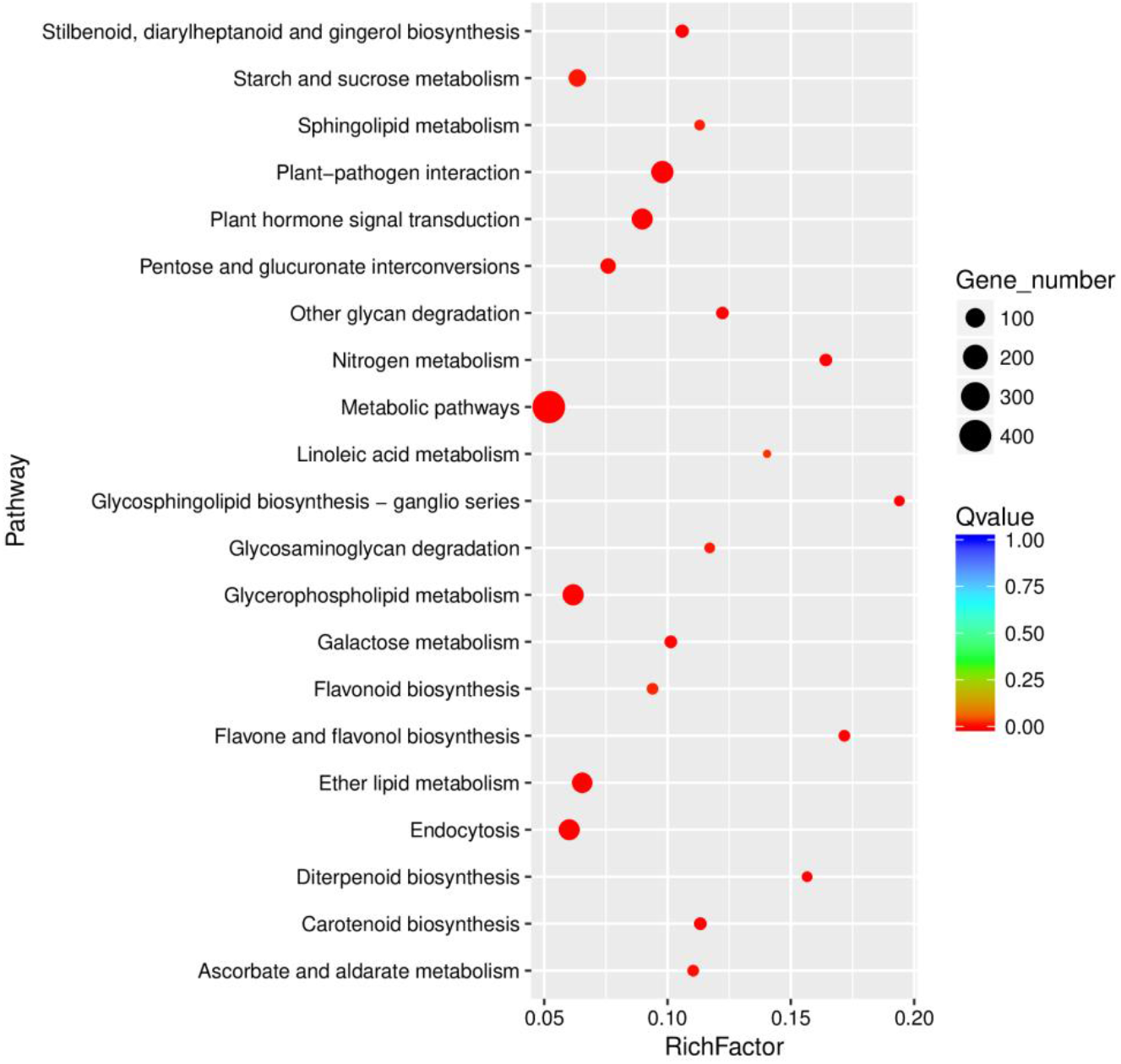
Top 20 KEGG pathways enriched in DEGs.

## DISCUSSION

### Identification of DEGs involved in chloroplast division

Leaf-color mutants are affected by chloroplast development and division, which can directly or indirectly affect chloroplast structure and number and other metabolic processes performed by these complex macromolecular machines, which have components positioned on both the inner and outer envelope surfaces(Gao et al. 2003). Many proteins, such as FtsZ1, tubulin-like protein, MinD, and ARTEMIS, are involved in chloroplast division(Yoshida et al. 2016;Fujiwara et al. 2019;Gonzalez-Garay et al. 1996;Dinkins et al. 2001;Funes et al. 2004). In this process, tubulin-like proteins localize to a ring at the site of plastid constriction, which drives chloroplast division. In our study, we found that six unigenes of seven DEGs encoding tubulin proteins were downregulated in the albino mutant, including Unigene5427_All, CL2099.Contig19_All, CL2099.Contig8_All, CL2099.Contig6_All, Unigene8230_All and CL2099.Contig4_All (Table S5), and one gene (Unigene1981_All) encoding an α-tubulin suppressor was upregulated, suggesting that these genes might be associated with chloroplast division and might therefore affect leaf color.

### Identification of DEGs involved in plant pigment synthesis

Plant leaf color is related to the types and contents of pigments in the cells. Enzymes are needed for the synthesis of plant pigments,; for example, Mg-chelatase and protoporphyrinogen oxidase are involved in chlorophyll synthesis, zeta-carotene desaturase and zeaxanthin epoxidase in carotenoid synthesis, and chalcone synthase in anthocyanin synthesis.

In our work, one gene (CL11028.Contig1_All) encoding magnesium chelatase was significantly downregulated, while Unigene17322_All, encoding protoporphyrinogen oxidase, was upregulated, as was one gene encoding the probable chlorophyll(ide) b reductase NYC1. Magnesium chelatase is a heterotrimeric enzyme complex that catalyzes a key regulatory and enzymatic reaction in chlorophyll biosynthesis(Rissler et al. 2002), protoporphyrinogen oxidase is a common enzyme in chlorophyll biosynthesis (Molina et al. 2010), and chlorophyll b reductase NYC1 and NOL (NYC-like) form the chlorophyll b reductase complex, which is necessary to catalyze the first step of chlorophyll b degradation(Sato et al. 2010).

Seven zeta-carotene desaturase genes (three upregulated and four downregulated), three zeaxanthin epoxidase genes (one upregulated and two downregulated), and one 15-cis-zeta-carotene isomerase gene (upregulated) were differentially expressed. Zeta-carotene desaturase is a key enzyme that controls beta-carotene production upstream of carotenoid biosynthesis(Wong et al. 2004). Zeaxanthin epoxidase is an important enzyme that converts zeaxanthin to violaxanthin(Thompson et al. 2000).15-Cis-zeta-carotene isomerase can isomerize 9,15,9’-tri-cis-zeta-carotene into 9,9’-di-cis-zeta-carotene, which is the last identified gene in the plant carotenoid biosynthetic pathway(Zambrano et al. 2015).

Five genes predicted to encode naringenin-chalcone synthase were downregulated, and Unigene26705_All was significantly decreased (|log_2_Ratio|>10). Chalcone synthase is the first key enzyme and rate-limiting enzyme in flavonoid synthesi(Ferrer et al. 2014).

Four genes encoding flavonoid 3’-monooxygenase were downregulated in the albino mutant. Flavonoid 3’-monooxygenase is responsible for one reaction in flavonoid compound biosynthesis(Larson et al. 1986).

We identified fifteen differentially expressed cytochrome P450 genes, of which three genes were upregulated and twelve downregulated. Cytochrome P450 is a monooxygenase encoded by a supergene family that is involved in the synthesis of many terpenoids in plants, including carotenoids(Marc et al. 2007). In addition, we found six genes encoding chloroplast chlorophyll a/b-binding protein, all of which were downregulated. Chlorophyll a/b-binding protein can enhance photosynthesis in the chloroplasts of plants(Lamppa et al. 1988).

### Identification of DEGs involved in anthocyanin transport

In our study, twenty-two genes encoding ABC (ATP binding cassette) transporter, five genes encoding MATE (multidrug and toxic compound extrusion) and eight genes encoding GST (glutathione S-transferase) were found among the DEGs. All three of these transporter types can transport anthocyanin in plants by various pathways. In this work, four genes encoding ABC transporters were upregulated in HUA, while the others were downregulated. The upregulated genes were CL5908.Contig1_All, Unigene14496_All, CL1076.Contig2_All and CL9731.Contig1_All, as shown in Table S5. Both the genes encoding MATE and those encoding GST were downregulated in HUA compared to LV. These results suggested that the transporters might transport plant pigments by complex molecular mechanisms.

### Identification of DEGs encoding transcription factors involved in leaf color

Transcription factors are involved in chloroplast development through upstream regulation of gene expression pertaining to chlorophyll, anthocyanin and carotenoid biosynthesis in plants, such as the MYB, bHLH and WD40 transcription factor families. MYB transcription factors, which regulate the expression of key enzymes in the early flavonoid biosynthetic pathway, enhance the synthesis and accumulation of proanthocyanidins in plants(Rosany et al. 2018). In tobacco, overexpression of *LeAN2*, which encodes an anthocyanin-associated R2R3-MYB transcription factor, induced the expression of several anthocyanin biosynthetic genes, and the content of anthocyanin was markedly higher in the transgenic tobacco than in wild-type plants[10]. Here, thirteen MYB transcription factors were identified; one was upregulated and the others downregulated. Recent studies have revealed that bHLH transcription factors also regulate chlorophyll biosynthesis. For example, PIF1 negatively regulates chlorophyll biosynthesis, while *pif1* mutant seedlings accumulate excess free protochlorophyllide when grown in the dark, with consequent lethal bleaching upon exposure to light(Huq et al. 2004). In this study, we found ten bHLH family genes that were downregulated in HUA, as shown in Table S5. WD40 transcription factors regulate the synthesis and accumulation of anthocyanin by affecting the expression of anthocyanin structural genes(Carey et al. 2004). TTG1, a WD-repeat (WDR) protein, acts as an important regulator of enzymes controlling proanthocyanidin biosynthesis in plants (Baudry et al. 2004). In the present study, five genes encoding WD40 were downregulated, while one was upregulated. These results suggested that transcription factors might constitute a complex network regulating leaf color in plants.

## ACKNOWLEDGMENTS

This study was supported by the National Natural Science Foundation of China (32002083), Henan Science and Technology Research Project (172102110248, 172102110151, and 172102110263), and Open Fund of Henan Key Laboratory of Tea Comprehensive utilization in South Henan (HNKLTOF2018007).

